# Ebola virus infection of Flt3-dependent, conventional dendritic cells and antigen cross-presentation leads to high levels of T-cell activation

**DOI:** 10.1101/2024.05.14.594125

**Authors:** Linda Niemetz, Bianca S. Bodmer, Catherine Olal, Beatriz Escudero-Pérez, Katharina Hoehn, András Bencsik, Molly A. Vickers, Estefanía Rodríguez, Lisa Oestereich, Thomas Hoenen, César Muñoz-Fontela

## Abstract

Severe Ebola virus disease (EVD) is characterized by excess, dysregulated T-cell activation and high levels of inflammation. Previous studies have described that in vitro EBOV infection of monocyte-derived DCs (moDCs) inhibits DC maturation resulting in suppression of T-cell activation. However, it is unknown how other DC subsets distinct from moDCs respond to EBOV infection.

To better understand how DCs initiate T cell activation during EBOV infection, we assessed the response of FMS-like tyrosine kinase 3 (Flt3)-dependent, conventional mouse DCs (cDCs) to EBOV infection, and developed a DC-T-cell co-culture system utilizing a recombinant EBOV expressing the model antigen ovalbumin.

Our findings suggested that, in contrast to moDCs, cDC2 and cDC1 were poorly infected with EBOV, although both infected and bystander cDCs displayed high levels of activation. DCs were able to activate CD8 T cells via cross-presentation of EBOV antigens obtained from cell debris of EBOV-infected cells. Of note, rather than interfering with cross-presentation, EBOV infection enhanced the cross-presentation capacity of DCs.

Our findings indicate that EBOV infection of Flt3-dependent cDCs, results in activation rather than inhibition leading to high levels of CD8 T-cell activation. With that we propose a mechanistic explanation for the excess T-cell activation observed in severe human EVD.

## BACKGROUND

Ebola virus disease (EVD) is caused by infection with members of the *Orthoebolavirus* genus, often Ebola virus (EBOV, species *Orthoebolavirus zairense*). The severity of EVD has been linked to high inflammation, virus dissemination and excessive T-cell activation followed by lymphopenia ^1–5^. A robust but regulated T-cell response, is associated with EVD recovery ^4–8^.

The transition between innate and adaptive immunity is a key checkpoint in EBOV infection. Activation of CD8 T cells requires antigen-presentation in the context of major histocompatibility complex class I molecules (MHC-I) by professional antigen-presenting cells (APCs). Some APCs, especially dendritic cells (DCs), cannot only present intracellular antigens on MHC-I, but are able to also direct exogenously-acquired antigens to MHC-I in a process called cross-presentation. This strategy allows CD8 T-cell activation in the absence of direct virus infection of APCs ^9,10^.

DCs form a heterogenous class of immune cells ^11–14^ that develop in the bone marrow from common DC progenitors by stimulation with FMS-like tyrosine kinase-3 (Flt3) ^15^. This developmental pathway gives rise to plasmacytoid DCs (pDCs) ^16^ and conventional DCs (cDCs) ^11–14^. The latter can be further subclassified into two main subsets – cDC1 and cDC2 ^11–14^. Additional subclassifications of cDC1 and cDC2 that are shaped by the microenvironment are the object of ongoing research ^14,17–19^. Moreover, DCs can also derive from monocytes (moDCs) in inflammatory settings.

DCs have been identified as early infection targets of EBOV ^7,20^. However, previous studies have described that, in cell culture, EBOV inhibits moDC maturation and IFN-I signaling in infected cells, likely due to the IFN-I antagonist activity of two viral proteins, namely, viral protein 35 (VP35) and VP24. These findings, using moDCs as a model APC, led to the hypothesis that lack of DC maturation upon EBOV infection resulted in poor T-cell activation ^21–25^. However, the effect of EBOV infection on cDCs has not yet been explored.

In the present study, we investigated the response of different DC subsets to EBOV infection. We examined whether EBOV-infected mixed-DC cultures activated virus-specific CD8 T cells, and dissected the role of different DC subsets in T-cell activation. To do so, we generated a novel recombinant EBOV expressing the model antigen ovalbumin (OVA) as a non-structural protein. Our findings indicated that EBOV preferentially infected moDCs and that pDCs were spared from infection. Contrary to moDCs, cDCs were highly activated upon EBOV infection. Futhermore, cDC were able to cross-present EBOV antigens to CD8 T cells which was further enhanced by EBOV infection. Our findings provide a mechanistic explanation for the excess T-cell activation observed in severe human EVD.

## METHODS

### Mice

C57Bl/6 mice were purchased from Jackson Laboratories and Thy1.1xOT1xC57Bl/6 mice were obtained from the Helmholtz Center for Infection Research Braunschweig, Germany. Both strains were bred in the animal facility at the Bernhard Nocht Institute for Tropical Medicine in Hamburg (BNITM). All mice were euthanized for organ collection at 9-15 weeks of age, which was approved by German animal welfare authorities (*Behörde für Gesundheit und Verbraucherschutz*, Approval 2021-T007).

### Generation of bone marrow-derived DCs

Bone marrow cells were isolated from femurs and tibiae of mice and cultured (2×10^6^ cells/mL, 6-well culture plates) for 8 days at 37°C and 5% CO2 in RPMI1640 (Life technologies) supplemented with 10% FBS, 500ng/mL Penicillin/ Streptomycin, 100ng/mL (Flt3L) (Peprotech), and 2mM beta-mercapthoethanol. On day 3, half of the medium was replaced with medium containing fresh Flt3L. Non-adherent cells were harvested at day 8.

### OT-1 CD8 T-cell isolation

OT-1 CD8 T cells were isolated from Thy1.1xOT-1 mice spleen by negative selection (Miltenyi Biotec).

### Generation of recombinant EBOV-OVA

The OVA ORF was cloned as an additional transcriptional unit between the NP and VP35 genes into a cDNA plasmid encoding Zaire ebolavirus rec/COD/1976/Mayinga-rgEBOV (GenBank accession number KF827427.1, rgEBOV)^26^ doubling the NP/VP35 non-coding region. Rescue of the recombinant virus was performed as previously described^27^ in the BSL4 laboratory at the FLI.

### Infections

All experimental infections were performed within the BSL4 facility at the BNITM. Cells were infected with EBOV H.sapiens-tc/COD/1976/Yambuku-Mayinga (wildtype (wt)EBOV) or recombinant EBOV-OVA. DCs were infected with MOI 3 and VeroE6 cells with MOI 0.1.

### Harvest of cell debris and extracellular vesicles (EVs)

VeroE6 cells were infected with recombinant EBOV-OVA (MOI 0.1). After 7 days, infected cells were inactivated by UV irradiation (wavelength 254nm, distance 15cm fisher scientific REF: 10712467) for 2x 10 min. Absence of infectious viral particles was confirmed by immunofocus assay. Inactivated cell culture supernatant was collected and cell debris was pelleted by centrifugation at 2000xg for 10min. Supernatant was discarded and the cell debris pellet was resuspended in RPMI 1640 media, supplemented with 10% FBS. For the collection of EVs, VeroE6 cells were infected with EBOV-OVA in a T75 cell culture flask (MOI 0.1). After 5 days the cell culture media was exchanged to EV-depleted DMEM supplemented with 2.5% FBS and incubated for another 2 days. Media was depleted of EVs by ultracentrifugation at 100,000xg for 18 hours at 4°C. EVs were isolated from cell culture supernatant by differential centrifugation (500xg and 2000xg), ultrafiltration (Amicon filter 10kDA) and size-exclusion chromatography (IZON qEV Gen2 columns).

### DC-T cell co-culture

DCs were infected with wt EBOV, EBOV-OVA (MOI 3) or were mock-infected. Twenty-four hours post-infection, DCs (3×10^5^ cells/well, 96-well plate) were stimulated with cell debris (from one well of 6-well plate per reaction, 100µL) or EV (from one T75 flask per reaction, 100µL) from EBOV-OVA infected or mock-infected VeroE6 cells or media control (non-stimulated control) for 3-5 h. Uninfected DCs were stimulated with 100µg/mL soluble ovalbumin (sOVA). DCs were then co-cultured with CellTrace Violet stained OT-1 CD8 T cells (500,000 cells/well) for 4 days.

### Flow Cytometry and FACS

Cells were harvested and Fc-receptors were blocked with CD16/CD32 Fc-Block antibody. Cells were stained with Zombie NIR (BioLegend) for 15 min, followed by staining with an antibody cocktail for 30 min. All fluorochrome-conjugated antibodies were purchased from BD Biosciences or BioLegend. EBOV-OVA infection was determined with anti-MHC-I-SIINFEKL (APC, 25-D1.16) and anti-EBOV-GP (5D2, 5E6; AF555). Monoclonal antibodies against EBOV-GP (5D2 and 5E6) have been previously described.^7,28^ Samples were inactivated with Cytofix/Cytoperm (BD Biosciences) containing 4% formaldehyde for 60 min and were acquired using a Cytek Aurora and analyzed with FlowJo Software. Cells were sorted with a FACS Aria III instrument.

### Statistical analysis

Statistics were performed using GraphPad Prism 10 software. Mock-infected and EBOV-infected groups were compared by non-parametrical t-test for unpaired samples (Mann-Whitney test).

## RESULTS

### EBOV preferentially infects moDCs and induces cDC activation

We performed experimental infections of mixed DC cultures, containing moDCs, cDC1, cDC2 and pDCs, generated by stimulating mouse bone marrow progenitor cells with Flt3-ligand (Flt3L) ^29^. After differentiation, 91% of live cells expressed the DC marker CD11c, of which on average 18% were identified as cDC1 (CD24^+^/ XCR1^+^/CD11b^-^/SIRPa^-^), 10% as cDC2 (CD24^-^/XCR1^-^/CD11b^+^/SIRPa^+^), 19% as pDCs (CD24^-^/XCR1^-^/CD11b^-^/SIRPa^-^/SiglecH^+^/Ly6C^+^) and 0.6% as moDCs (CD14^+^/F4/80^+^/CD64^+^/CD11b^+^) (Fig. 1A, Supplementary Fig. 1). MoDCs showed high expression levels of the activation markers CD86 and CD80 in the steady state. While cDCs were highly activated upon stimulation with lipopolysaccharide (LPS), pDCs were unresponsive to LPS stimulation (Fig. 1C).

**Figure 1.**
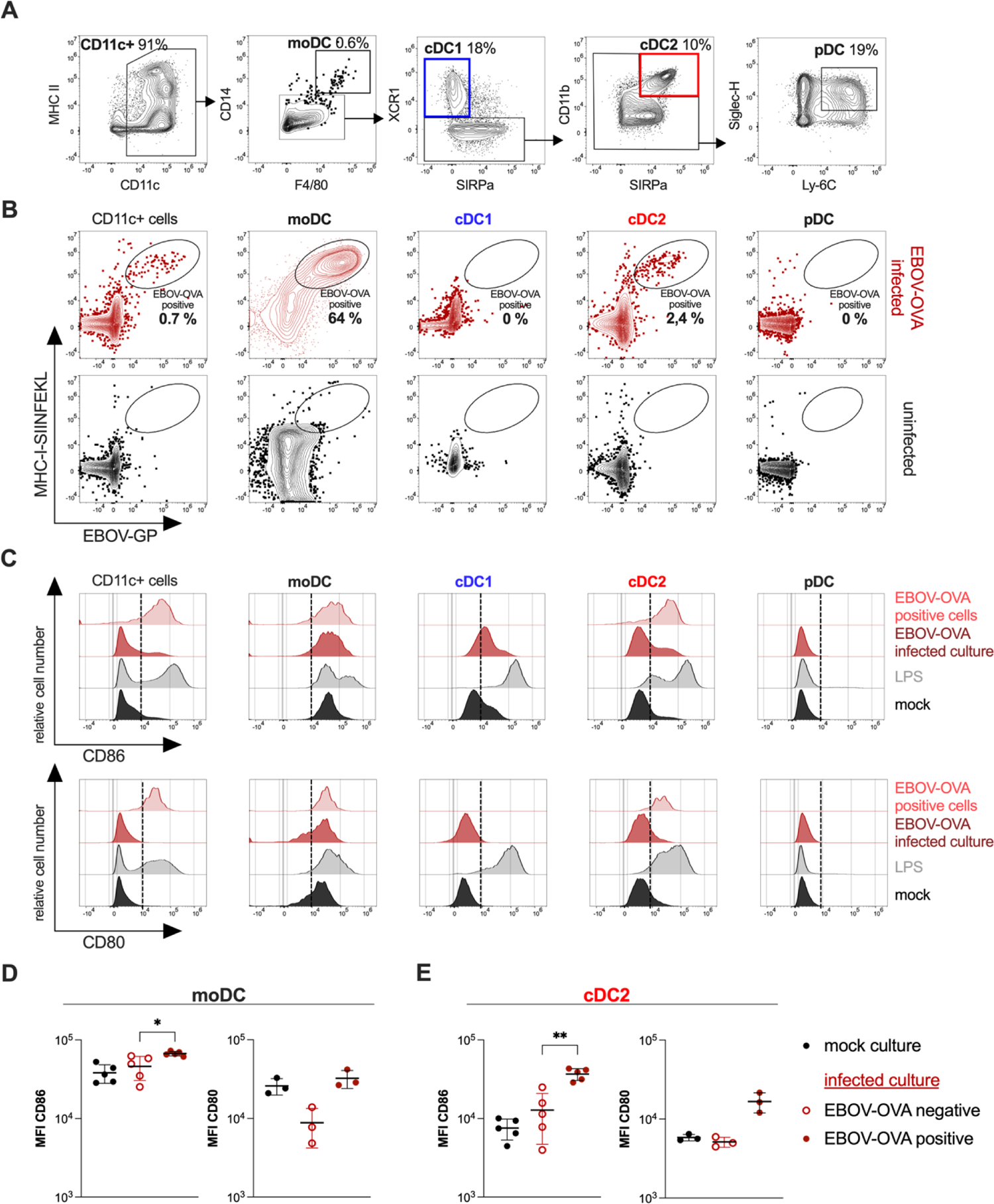
Characterization and EBOV-OVA infection of a mixed DC culture. DCs, derived from bone marrow progenitor cells supplemented with Flt3L for 8 days, were infected with EBOV-OVA (MOI=3) for 24 hours. **(A)** DC culture consisted of moDC*s*, cDC1, cDC2 and pDC*s* as characterized by flow cytometry. **(B)** EBOV-OVA positive cells in all CD11c^+^ cells, or separate subsets in EBOV-OVA infected (red) or mock-infected DC culture (black) were identified by the surface expression of EBOV glycoprotein (EBOV-GP) and MHC-I-SIINFEKL. Population frequencies (percentage of DC subset population) reflect mean value of 5 independent experiments (individual values given in Figure S1). **(C)** CD86 and CD80 surface expression *on* all CD11c^+^ cells, or on separate DC subsets in mock-infected – (black), LPS-stimulated – (grey), or EBOV-OVA infected DC culture (dark red), and *on* EBOV*-*OVA positive cells within *the* infected culture (light red). Representative plots of n=5 (CD86) or n*=*3 (CD80) independent experiments. (D) Mean fluorescence intensity (MFI) of CD86 (n=5) and CD80 (n=3) in EBOV-OVA negative - (red, empty) and EBOV-OVA positive (red, filled) moDCs and cDC2 compared to mock*-*infected moDCs and cDC2 (black). Non-parametric t-test (Mann-Whitney test) *<0.05, **<0.01.

We reasoned that, with the exception of resident macrophages, this mixed DC culture recapitulated the main APCs located at the natural portals of EBOV entry. Thus, we could use the mixed DC culture as a model to explore EBOV infection of APCs at early stages of infection. With this goal, we utilized a recombinant EBOV-OVA which expressed OVA at high levels in infected cells (Supplementary Fig. 2).

We reasoned that infected cells could be identified by surface expression of the EBOV-glycoprotein (GP) as a consequence of viral budding ^7^. In addition, previous studies have shown that OVA expressed in cells is cleaved into the immunodominant, H-2K^b^-restricted, CD8 T-cell peptide SIINFEKL and that MHC-I-SIINFEKL complexes can be detected by the specific monoclonal antibody 25-D1.16 ^30^. Thus, we utilized this antibody to identify EBOV-OVA infected cells and cells presenting viral antigens. Specific double staining of EBOV-GP and MHC-I-SIINFEKL in EBOV-OVA infected DC cultures revealed that less than 1% of CD11c^+^ cells were infected at 24 hours post-infection.

Despite being an underrepresented myeloid population in this setup, moDCs showed the highest infection rate of 64%. Within the cDC2 population 1-4% were EBOV-OVA positive, while cDC1 and pDCs were spared from infection. MHC-I-SIINFEKL was detected on the surface of EBOV-GP-positive moDCs and cDC2, suggesting that viral antigens were presented in the context of MHC-I by EBOV-infected DCs. In bystander (non-infected) DCs, MHC-I-SIINFEKL could not be detected (Fig 1B and Supplementary Fig. 3).

Next, we wanted to assess the effect of EBOV infection on the activation status of infected and bystander DCs. Moderate differences in the expression levels of CD86 and CD80 were found between infected and non-infected moDCs. However, infection of cDC2 resulted in significant increase of expression levels of both CD86 and CD80 indicating that this population responded to infection with activation. Interestingly, uninfected cDC1, as well as some uninfected cDC2, showed increased expression of CD86 but not CD80, suggesting partial activation, which was not observed in pDCs (Fig. 1C, D and Supplementary Fig. 4).

To summarize, in a mixed DC culture moDCs and, to some extent cDC2, were preferred infection targets of EBOV, and infected cells presented viral antigens in the context of MHC-I. As opposed to moDCs, infected as well as bystander cDC1 and cDC2 showed a highly activated phenotype. We next wanted to address the consequences of these activation profiles for T-cell immunity.

### EBOV-OVA infected DCs activate cognate CD8 T cells

Previous studies indicated that EBOV-infection of moDCs inhibited the capacity of these cells to induce T-cell activation ^21–25^. Since we observed that, cDCs showed signs of activation, we wanted to determine whether these mixed DCs could activate cognate CD8 T cells. For that, we utilized CD8 T cells isolated from OT-1 mice, a mouse strain with a transgenic TCR designed to recognize MHC-I-SIINFEKL complexes ^31^. Thus, we co-cultured EBOV-OVA infected mixed DC cultures with CellTrace Violet-stained OT-1 CD8 T cells 24 hours post-infection and analyzed T-cell proliferation and activation by flow cytometry after 4 days (Fig. 2A). We examined T-cell proliferation by the signal reduction of the proliferation dye CellTrace Violet as it is progressively diluted with every generation of cell division. EBOV-OVA infected mixed DC cultures induced strong T-cell proliferation as opposed to mock-infected DCs (Fig. 2B). Moreover, proliferating OT-1 CD8 T cells were highly activated compared to steady state OT-1 CD8 T cells, as shown by upregulation of the surface activation markers CD44, CD69 and CD25, and downregulation of CD62L (Fig. 2C).

**Figure 2.**
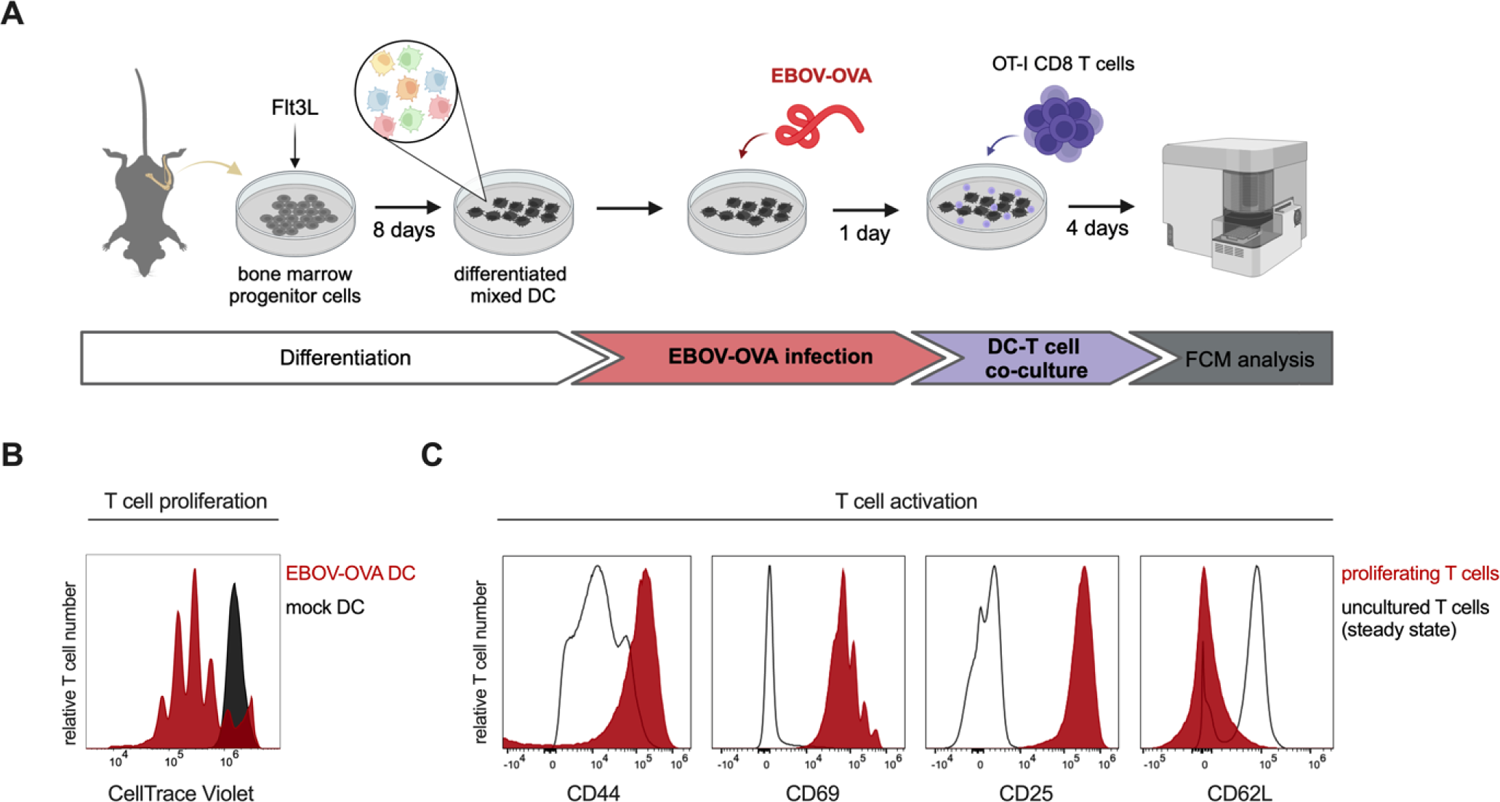
T-cell proliferation and activation by EBOV-OVA-infected DCs. **(A)** Experimental setup: DCs were derived from bone marrow progenitor cells, incubated with Flt3L for 8 days, infected with EBOV-OVA for 24 hours and co-cultured with CellTrace Violet stained OT-1 CD8 T cells for four days. T cells were analyzed by flow cytometry (FCM). **(B)** T-cell proliferation upon co-culture with EBOV-OVA*-*infected (red) or uninfected DCs (black). T*-*cell proliferation was assessed as *a* reduction of the CellTrace Violet signal intensity as a consequence of cell division. **(C)** T-cell surface activation markers CD44, CD69, CD25, CD62L in proliferating T cells (red) compared to steady state T cells before co-culture with DC (black line). Representative plots of 3 independent experiments.

Taken together, our data indicated that a mixed DC culture, containing EBOV-infected and bystander moDCs, cDCs and pDCs, was able to activate cognate CD8 T cells. Next, we wanted to dissect the specific contribution of each DC subset to T-cell activation.

### Bystander DCs activate CD8 T cells via cross-presentation

Direct infection of DCs is not required for antigen presentation in the context of MHC-I, because DCs bear the capacity to direct exogenously acquired antigens to MHC-I in a process called cross-presentation. Thus, not only EBOV-OVA infected DCs may be able to activate OT-1 CD8 T cells, but also bystander DCs that acquired antigens released by infected-cells. Due to the fact that EBOV infected less than 1% of the mixed DCs, we next wanted to investigate whether bystander DCs were able to activate EBOV-specific T cells via cross-presentation.

Previous studies have shown that extracellular vesicles (EVs) from EBOV-infected cells may contain viral proteins ^32,33^. Furthermore, infected cells may also transfer viral antigens to bystander DCs via cell debris caused by cell death. Therefore, we wanted to test whether EVs or cell debris from EBOV-OVA-infected epithelial cells could serve as antigenic source for cross-presentation.

We stimulated mixed DC cultures with EVs or cell debris from EBOV-OVA infected VeroE6 cells (Fig. 3A, Supplementary Fig. 5). Since VeroE6 cells are of monkey origin, they do not express mouse MHC-I and therefore cannot directly present antigens to OT-1 T cells. Importantly, to avoid direct infection of DCs by EBOV-OVA carried along the EV and cell-debris isolation procedures, we inactivated EVs and cell debris samples by UV irradiation before using them for DC stimulation.

**Figure 3.**
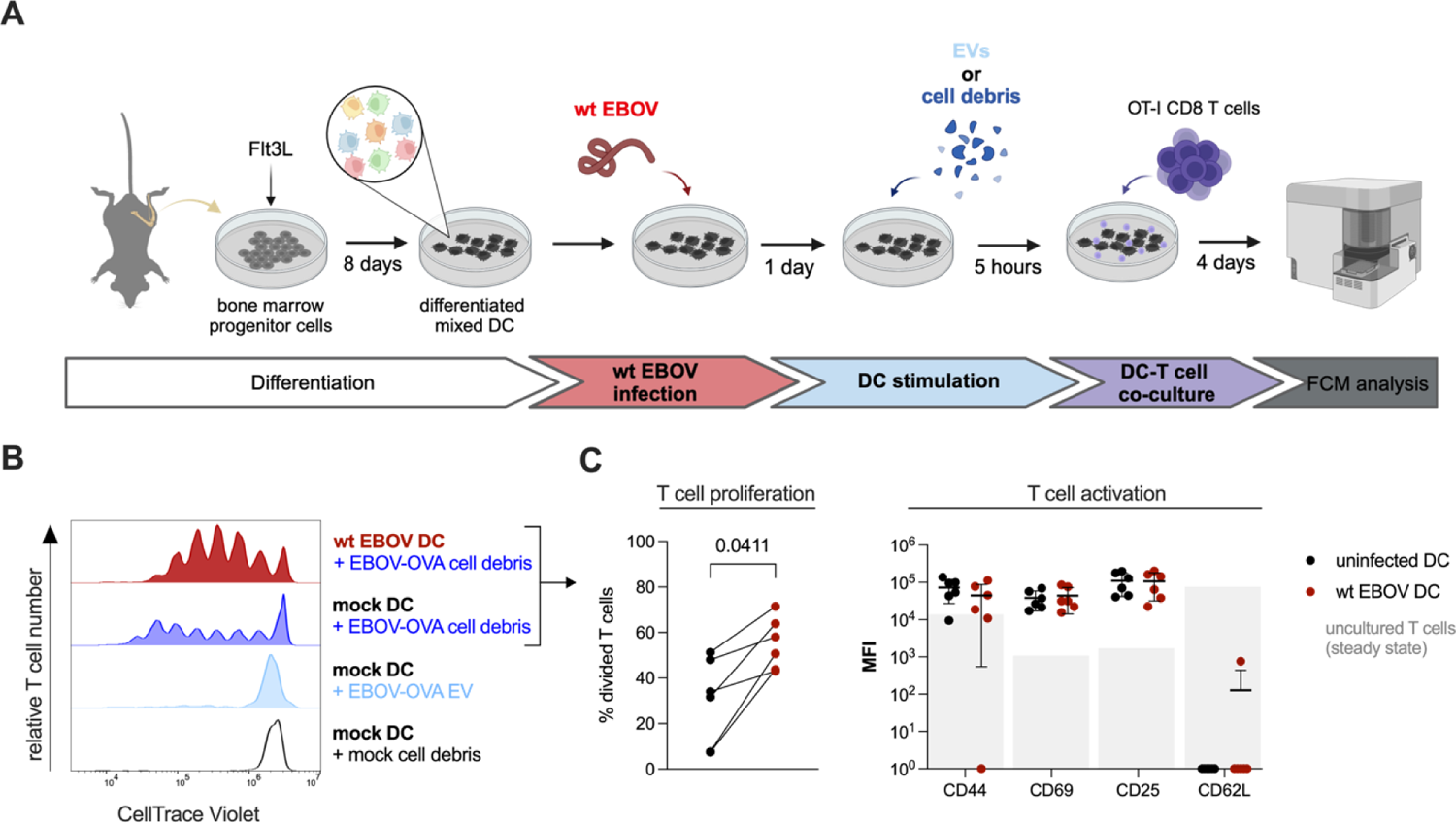
T-cell proliferation upon cross-presentation of cell debris or EVs by uninfected or wt EBOV-infected DCs. **(A)** Experimental setup: Mixed DC cultures were infected with wt EBOV (red) for 24 hours or left uninfected (black) and then stimulated with cell debris or EVs. Cell debris was collected from uninfected (mock cell debris) or EBOV-OVA-infected epithelial cells (EBOV-OVA cell debris). EVs were collected from EBOV-OVA-infected epithelial cells (EBOV-OVA EV). Stimulated DCs were co-cultured with OT-1 CD8 T cells for four days. T cells were analyzed by flow cytometry (FCM). **(B)** T-cell proliferation induced by uninfected DCs stimulated with mock cell debris (black), EBOV-OVA cell debris (dark blue) or EBOV-OVA EV (light blue), and T-cell proliferation induced by wt EBOV infected DCs stimulated with EBOV-OVA cell debris (red). Peak height indicates the number of T cells. **(C)** Percent of T cells driven into proliferation upon co-culture with uninfected DCs (black) or wt EBOV infected DCs (red) stimulated with EBOV-OVA cell debris. And, respective median fluorescence intensity (MFI) of T-cell activation markers CD44, CD69, CD25, CD62L in proliferated T cells compared to uncultured, steady state T cells (grey bars). Lines connect pairs from the same mouse donor (biological pairs). (n=6, non-parametric unpaired t-test, Mann-Whitney test).

Our results showed that DCs stimulated with cell debris from EBOV-OVA-infected donor cells activated OT-1 CD8 T cells and induced T-cell proliferation. EV-mediated DC stimulation, on the other hand, did not result in CD8 T-cell proliferation, suggesting that cell debris as opposed to EVs was the main source of EBOV antigen for cross-presentation (Fig. 3B). These results strongly suggested that bystander DCs in a mixed DC culture contributed substantially to the activation of EBOV-specific T cells via cross-presentation.

### EBOV infection enhances cross-presentation

Antigen cross-presentation is a highly regulated process, not only from the host perspective to avoid T-cell mediated immunopathology, but also some viruses have evolved mechanisms to interfere with cross-presentation in order to avoid the host CD8 T-cell response ^34–36^. Therefore, we next wanted to test the effect of EBOV infection in DC-mediated cross-presentation.

We infected mixed DC cultures with wt EBOV 24h prior to stimulation with UV-inactivated cell debris from EBOV-OVA-infected VeroE6 cells. DCs infected with wt EBOV cannot induce OT-1 CD8 T-cell proliferation (Supplementary Fig. 6). Thus, by comparing wt EBOV infected vs mock-infected DCs, we could assess the effect of infection in the cross-presentation capacity of DCs.

Similarly to mock-infected DCs, wt EBOV-infected DCs induced T-cell proliferation and activation via cross-presentation of antigens from cell debris obtained from EBOV-OVA infected VeroE6 cells. However, cross-presentation by wt EBOV-infected DC cultures resulted in higher T-cell numbers per generation (peak height per generation) and overall higher percentage of proliferating T cells (Fig. 3B). Proliferating T cells in both conditions displayed similar activation status as assessed by surface expression of T-cell activation markers (Fig. 3C and Supplementary Fig. 7).

Taken together, these results suggested that EBOV infection of DCs not only resulted in T-cell activation, but further increased the capacity of DCs to cross-present EBOV antigen to T cells. Next, we sought to determine the specific contribution of DC subsets to cross-presentation.

### cDC1 highly contribute to EBOV specific T-cell activation via cross-presentation

To dissect the roles of cDC1 and cDC2 in T-cell activation, we FACS-sorted cDC1 and cDC2. Next, we infected only the cDC2 culture with EBOV-OVA and thereafter, co-cultured infected cDC2 and uninfected cDC1 at different ratios. Twenty-four hours post infection, we added OT-1 CD8 T cells to the co-culture and analyzed T-cell proliferation and activation 4 days after (Fig. 4A).

**Figure 4.**
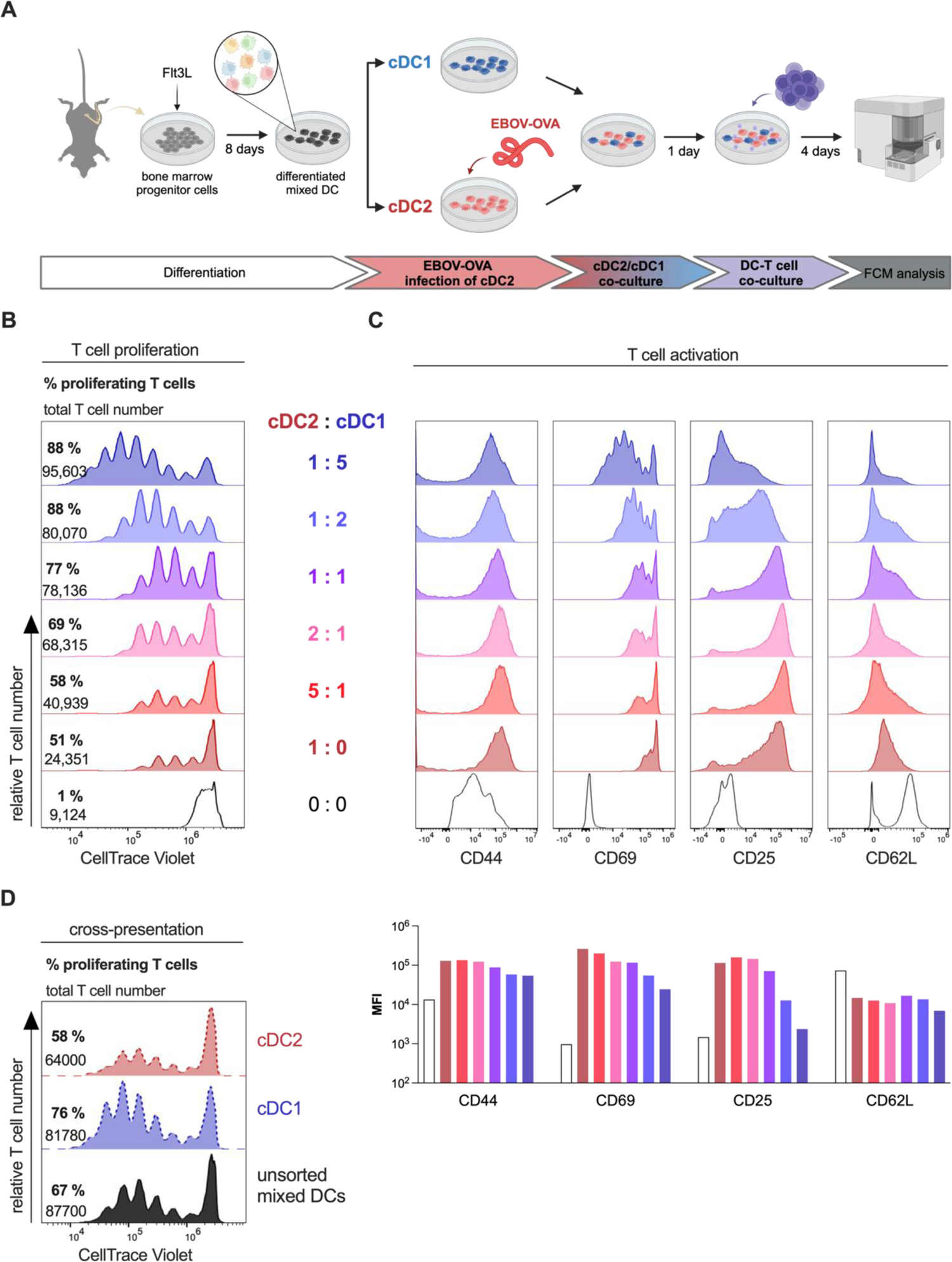
T cell proliferation and activation by cDC2/cDC1 co-cultures of varying ratios. **(A)** Experimental setup: cDC1 and cDC2 were FACS-sorted from mixed DC cultures. cDC2 were infected with EBOV-OVA (MOI=3) and subsequently co-cultured with uninfected cDC1 in different ratios. OT-1 CD8 T cells were added to the co-culture 24 hours post-infection and analyzed after 4 days. (**B**) T cell proliferation plots for different cDC2/cDC1 ratios. Numbers of cDC1 are increasing and numbers of cDC2 are decreasing from bottom to top. Numbers represent the total T-cell numbers and the percentage of proliferating T cells for each condition. **(C)** (top) Histograms of T-cell activation marker CD44, CD69, CD25 and CD62L expression in proliferating T cells compared to steady state, un-cultured T cells (black line). And (bottom), median fluorescence intensity (MFI) of T cell activation markers in different conditions. Data were acquired in a single experiment. (**D**) T-cell proliferation upon co-culture with unsorted DCs (black), cDC1 (blue) or cDC2 (red) stimulated with cell debris. Numbers represent the total T-cell number and percentage of proliferated T cells.

The EBOV-OVA infected cDC2 culture alone, induced T-cell proliferation and activation. However, T-cell proliferation was progressively augmented by increasing the numbers of cDC1 in the co-culture, as reflected by growing numbers of generations (peak number) and proliferating T-cell numbers (peak height). This suggested an important contribution of cDC1 to the observed T-cell activation (Fig. 4B). T cells showed similar expression of the T-cell activation markers CD44 and CD62L for all cDC2:cDC1 ratios. Conversely, CD69 and CD25 surface expression in proliferating T cells decreased with increasing numbers of cDC1 in the co-culture due to enhanced T-cell proliferation (Fig. 4C). In summary, bystander cDC1 played a key role in T-cell activation upon EBOV infection in a mixed DC culture, presumably via cross-presentation.

To test this hypothesis, we tested the cross-presentation capacity of each individual DC subset. We stimulated sorted cDC1s and cDC2s with cell debris from EBOV-OVA infected epithelial cells and co-cultured them with OT-1 CD8 T cells. While both DC subsets were able to cross-present exogenous antigens to some extent, cDC1 induced a stronger CD8 T-cell proliferation than cDC2, which suggested that, in the context of EBOV infection, cDC1s are the main cross-presenting DC population (Fig. 4D).

## DISCUSSION

In this study we wanted to determine the preference of EBOV for different DCs subsets and to evaluate the effect of infection on DC activation, antigen presentation and T-cell immunity. In this regard, despite its limitations, the mouse model has some advantages as follows: Firstly, despite being resistant to disease, mice experimentally infected with EBOV support high levels of virus replication and show evidence of virus dissemination ^7^. Moreover, EBOV infects mouse DCs in vivo and in vitro with virus titers similar to those observed in human DCs ^7,37^. Perhaps more importantly, the mouse model allowed us to track T-cell responses to a model antigen (OVA) expressed as a viral-encoded protein and to characterize DC-induced T-cell responses.

Previous studies have shown that EBOV infection of moDCs inhibited DC maturation resulting in poor T-cell activation ^21–25^. These results, however, were in conflict with the clinical picture of hyperinflammation and excessive T-cell activation observed in severe EVD ^1,4,5^. Previously, we observed that EBOV has preference for inflammatory DCs in mice, and seemed to spare other DC subsets ^7^. These findings raised the question of whether all DC subsets were functionally impaired upon EBOV infection or if certain DC subsets remained functional.

In an attempt to recapitulate the main APCs present in skin and mucosae, the main portals of EBOV entry ^38^, we derived a mixed-DC culture from bone marrow progenitor cells by stimulation with Flt3L. The mixed DC culture represented steady state cDC1, cDC2 and pDCs, ontogenetically distinct from moDCs^39–41^. However, our mixed DC culture also contained a small cell population that we characterized as moDCs due to the distinct surface expression of CD14, F4/80 and CD64. These markers, as well as SIRPa and CD11b, can be found in moDCs and various subclassifications of cDC2, making it difficult to distinguish between these subsets^17^. However, the population that we identified as moDCs in our study showed unique characteristics with regards to cell shape, basal activation marker expression, and infectability, which supported their distinction from immature cDC2 ^42^.

Despite their low percentage in the mixed DC culture, moDCs were the main target of EBOV, which strongly suggests that they may support high levels of virus replication at early stages post-infection. These results are also in agreement with the finding that high expression of Siglec-1 in activated moDCs may be a key attachment factor favoring entry of EBOV in these cells ^43^.

Besides moDCs, cDC2 were also susceptible to EBOV infection. A reason for that could be the phenotypical similarity between moDCs and cDC2 explained above. In contrast, cDC1 and pDCs were protected from EBOV infection. The resistance of cDC1s to infection with other enveloped viruses has been previously described, and has been linked to the distinct expression of the vesicle trafficking protein RAB15 as well as to langerin-TRIM5α-dependent virus recirculation to autophagosomes ^44,45^. Whether the same or similar molecular mechanisms protect cDC1 also from EBOV remains an open question. Like cDC1, pDC were not infected by EBOV, which is in agreement with previous studies utilizing EBOV-like particles ^46^. pDCs have been shown to be susceptible to some viruses, such as human respiratory syncytial virus, vesicular stomatitis virus or Sendai virus,^47,48^ but are refractory to most other viruses.

After EBOV infection moDCs only moderately increased the expression of CD86 and CD80, which was in line with previous findings demonstrating inhibition of moDC maturation upon EBOV infection by the action of VP35 and VP24 ^21–25^. In contrast, cDC2 were highly activated after EBOV infection. In fact, also bystander cDC2 and cDC1 within the infected mixed DC culture showed upregulation of the maturation marker CD86, suggesting partial DC activation as CD86 is shown to be upregulated earlier than CD80 during maturation ^49^. pDCs, on the other hand, showed no activation upon EBOV infection, which is in agreement with previous studies^46^.

Using recombinant EBOV-OVA, we showed that EBOV-antigens were loaded onto MHC-I and were presented by infected DCs. Furthermore, OVA-specific OT-1 CD8 T cells were activated and proliferated upon co-culture with EBOV-OVA-infected mixed-DC cultures. This result suggested that infected DCs were not only activated, but retained the ability to prime cognate T cells. These findings are also in agreement with the strong T-cell activation observed in EVD ^1,4,5^.

Bystander DCs were able to activate OT-1 CD8 T cells via cross-presentation of antigens for which, cell debris rather than EVs, was the main source. Our data indicated that cDC1 was the main cross-presenting DC population in the context of EBOV infection. This result is in agreement with previous work indicating that cDC1s are quite resistant to infection but highly specialized in antigen cross-presentation ^50^. We speculate that processing of EBOV antigen by cDCs at early stages of infection and migration to tissue-draining lymph nodes, may be a key checkpoint for early initiation of EBOV-adaptive immunity.

Strikingly, EBOV infection enhanced, rather than reduced, cross-presentation. This was an unexpected finding, as evidence indicated that viruses in general evolve mechanisms to prevent antigen presentation as well as cross-presentation ^34–36^. We speculate that this mechanism may contribute, at least to some extent, to the excess inflammation and T-cell exhaustion-like phenotype observed in severe EVD ^1,5^.

In summary, in this study we show that EBOV induces strong T-cell activation partially through DC infection but also through antigen cross-presentation mediated by cDC1. Our findings suggest that targeting this DC subset, for example via specific delivery of antigens, could serve as a vaccination strategy to enhance T-cell responses against EBOV.

## FOOTNOTES

### Author Contributions

LN contributed to study design, performed all experimental work and wrote the manuscript with CMF. BSB performed rescue of recombinant EBOV-OVA. CO, BEP, KH, AB and MAV performed work that contributed to experimental data obtained in the BSL4 laboratory. ER, LO and TH contributed to the study design and provided funding. CMF designed and coordinated the study. All authors edited the manuscript.

## Acknowledgements

We thank Arne Düsedau, Michelle Heung, Stephanie Wurr, Ludmilla Unrau and Sabrina Bockholt for excellent technical support during our study.

