## Supplementary Figure 1 for "Ebola virus infection of Flt3-dependent, conventional dendritic cells and antigen cross-presentation leads to high levels of T-cell activation"

**A**

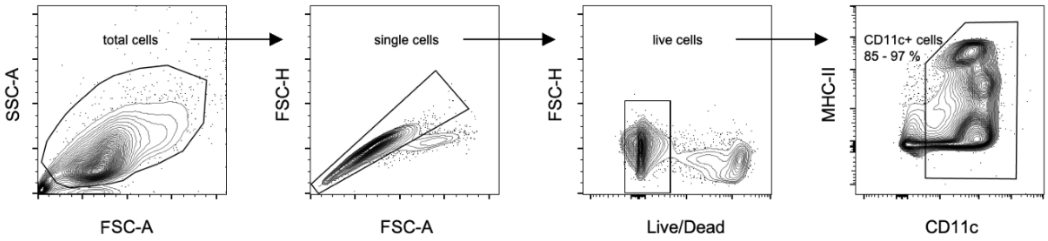


**B**

| | % of live cells | Frequency of DC subsets in % of CD11c+ | | | |
|---|---|---|---|---|---|
| replicates | **CD11c+ cells** | **moDC** | **cDC1** | **cDC2** | **pDC** |
| #1 | 97 | 0.3 | 34 | 16 | 20 |
| #2 | 89 | 0.5 | 27 | 16 | 17 |
| #3 | 87 | 0.5 | 10 | 7 | 20 |
| #4 | 91 | 1.4 | 12 | 6 | 18 |
| #5 | 93 | 0.5 | 8 | 4 | 19 |
| mean | 91 | 0.6 | 18 | 10 | 19 |

**C**

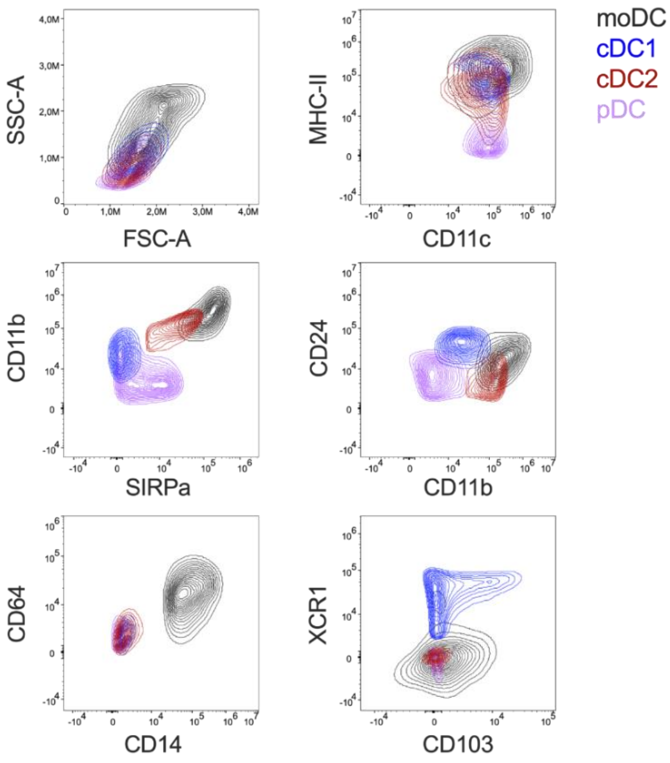


Niemetz et al. Supplementary Figure 1
