## Supplementary figures and images for "Ebola virus infection of Flt3-dependent, conventional dendritic cells and antigen cross-presentation leads to high levels of T-cell activation"

### Supplementary Figure 2

A

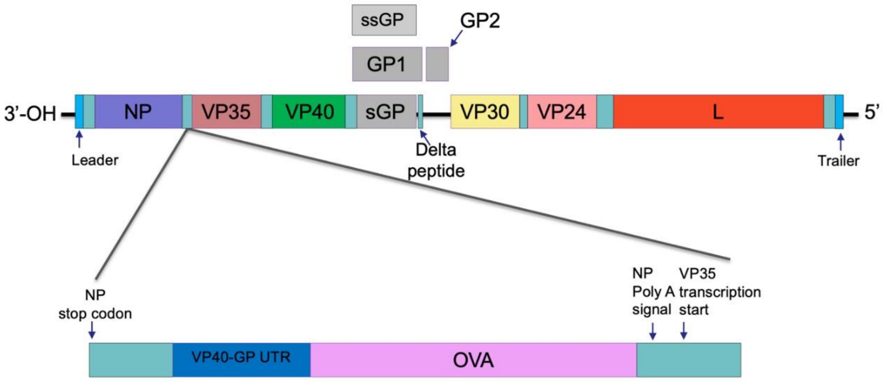


B

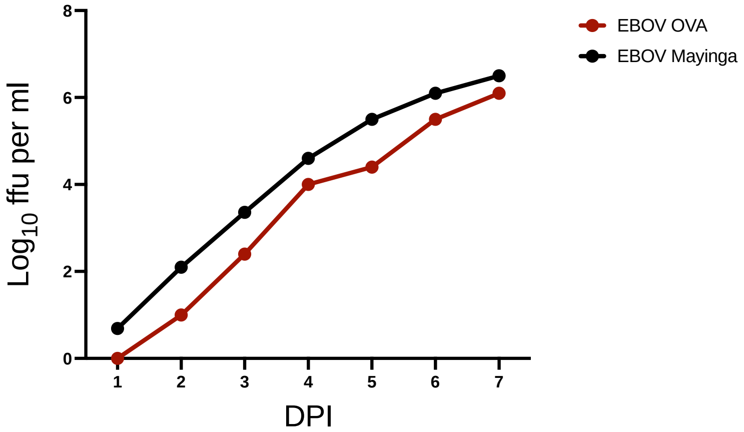


Niemetz et al. Supplementary Figure 2

### Supplementary Figure 4

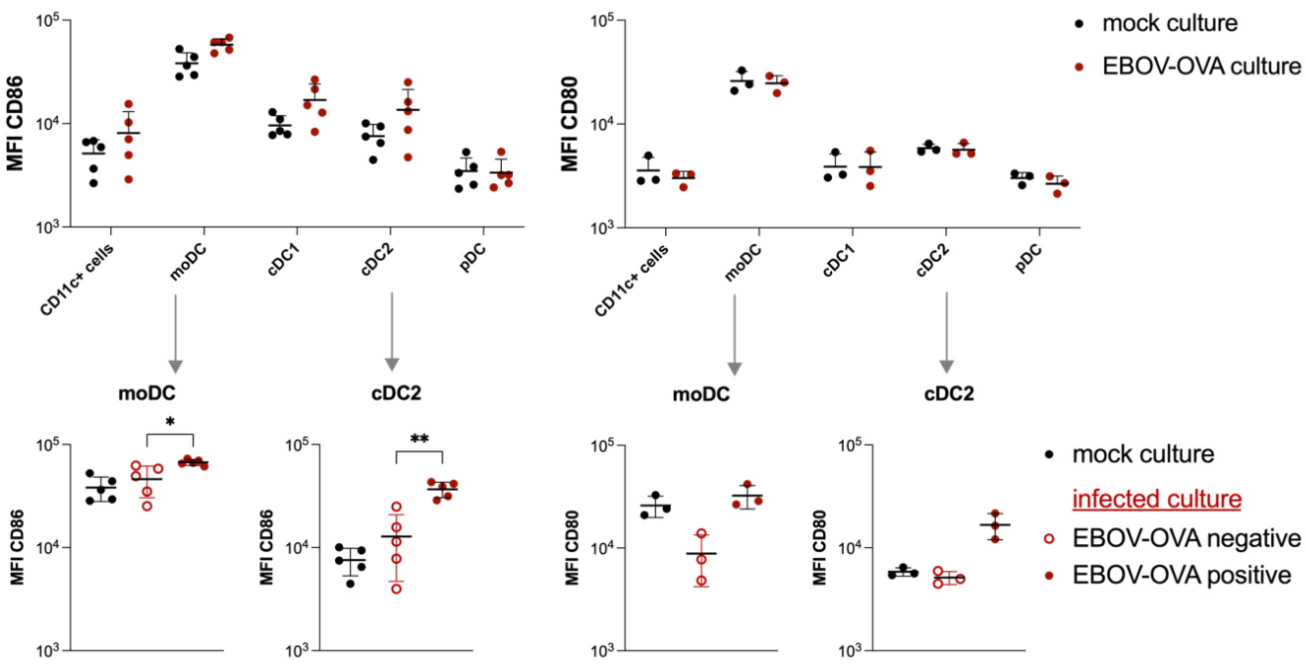


Niemetz et al. Supplementary Figure 4

### Supplementary Figure 5

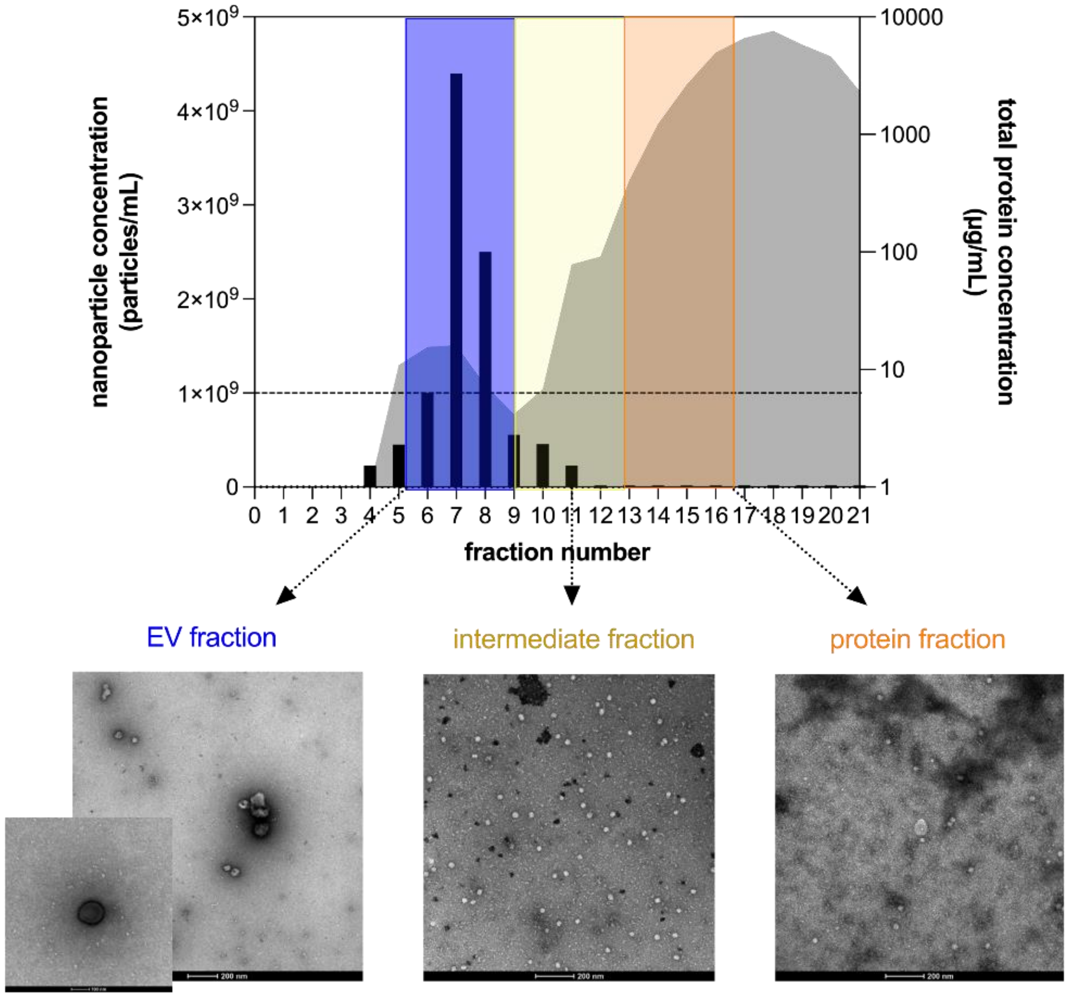


Niemetz et al. Supplementary Figure 5

### Supplementary Figure 6

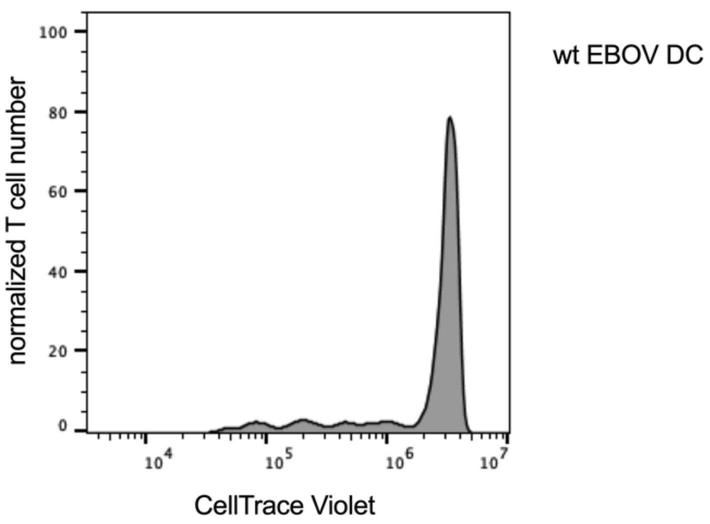

### Supplementary Figure 7

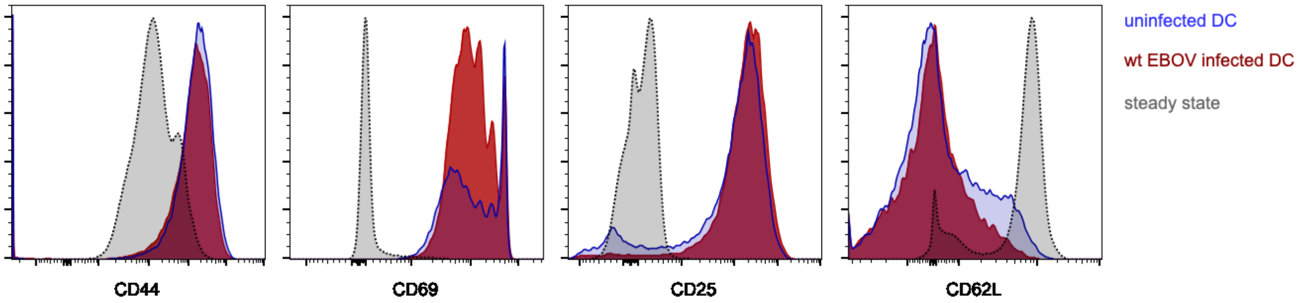


Niemetz et al. Supplementary Figure 7
