## Supplementary Figure 3 for "Ebola virus infection of Flt3-dependent, conventional dendritic cells and antigen cross-presentation leads to high levels of T-cell activation"

| condition | | Frequency of EBOV-OVA positive cells in % of DC subset: CD11c$^{+}$ cells | moDC | cDC1 | cDC2 | pDC | all other cells |
|---|---|---|---|---|---|---|---|
| MOI 3 | #1 | 0.54 | 71.9 | 0.01 | 0.73 | 0.01 | 0.09 |
| | #2 | 0.74 | 67.2 | 0.01 | 1.72 | 0.00 | 0.04 |
| | #3 | 0.94 | 81.9 | 0.01 | 4.96 | 0.02 | 0.03 |
| | #4 | 0.83 | 43.7 | 0.00 | 3.16 | 0.02 | 0.01 |
| | #5 | 0.42 | 55.2 | 0.01 | 1.61 | 0.04 | 0.02 |
| **mean** | | **0.70** | **64.0** | **0.01** | **2.44** | **0.02** | **0.04** |
| **MOI 3** | **LPS** | 0.93 | 35.1 | 0.00 | 1.01 | 0.01 | 0.00 |
| | **UV** | 0.02 | 0.26 | 0.00 | 0.03 | 0.01 | 0.00 |
| **MOI 1** | | 0.82 | 83.3 | 0.00 | 1.07 | 0.01 | 0.01 |

### = replicate number, LPS = stimulation of DCs with LPS 24 hours prior to infection, UV = UV-inactivation of EBOV-OVA prior to infection
